# Human-Perception-Aligned Machine Learning for Indoor–Outdoor Classification

**DOI:** 10.64898/2026.01.08.698168

**Authors:** Daniella Mahfoud, Zhiqun Tang, Raymond P. Najjar

**Affiliations:** Eye N’ Brain Research Group, Department of Ophthalmology, Yong Loo Lin School of Medicine, National University of Singapore, Singapore; Visual Neurosciences Group, Singapore Eye Research Institute, Singapore; Department of Biomedical Engineering, College of Design and Engineering, National University of Singapore, Singapore; Centre for Innovation & Precision Eye Health, Yong Loo Lin School of Medicine, National University of Singapore, Singapore; Ophthalmology and Visual Science Academic Clinical Program, Duke-NUS Medical School, Singapore

**Author notes:** Corresponding Author: Dr Raymond P. Najjar Department of Ophthalmology, Yong Loo Lin School of Medicine, National University of Singapore, Singapore 119228, Singapore.

**Keywords:** Human perception, outdoor light, machine learning, environmental light assessment, multispectral light data

## Abstract

Daylight is vital for eye, brain, and overall health, yet misperceptions of what constitutes true outdoor daylight exposure can lead to poor behavioural choices and weaken public health recommendations. We developed *InNOut*, a perception-aligned machine learning model trained on 73,879 minutes of multispectral light data to classify environments as indoor or outdoor, and benchmarked against public (n = 383) and expert (n = 17) judgments across 183 scenes. *InNOut* achieved excellent performance (AUC = 0.92 [95% CI, 0.92–0.93], sensitivity 73.9%, specificity 94.5%), closely aligning with expert (83.5%) and public (80.8%) judgments. Clear outdoor and indoor scenes were reliably classified, whereas ambiguous settings (e.g., windowed rooms, vehicles) were often judged indoor by participants but outdoor by the model, consistent with their spectral light profiles. *InNOut* bridges perception and measurement, offering a scalable means to map light environments and inform digital interventions in medicine and public health.

## INTRODUCTION

Light is one of the most powerful environmental cues influencing human physiology and behaviour. Across the lifespan, exposure to daylight supports ocular health, synchronizes circadian rhythms, and promotes psychological wellbeing.^1,2^ In children, greater time outdoors reduces myopia risk and slows its progression,^3,4^ with at least 120 minutes daily proving most effective.^5^ This protective effect is driven in part by daylight exposure itself, along with other factors such as richer spatial frequency content and reduced peripheral hyperopic defocus.^6,7^ Therefore, scalable interventions promoting time outdoors are on the rise for preventing or delaying the onset of myopia.^8,9^

Beyond ocular benefits, exposure to outdoor daylight supports broader cognitive and emotional development. In children, it enhances academic performance,^10^ sustains attention and reduces hyperactivity symptoms.^11^ In adults, more time outdoors is associated with better mood, lower stress, improved sleep, and overall psychological wellbeing.^12–15^ Daylight therapy and increased daylight exposure (*i.e.,* being outdoors for an additional 1-2 hours per day) have shown efficacy in regulating circadian rhythms and are associated with lower depressive outcomes,^16,17^ whereas insufficient exposure is linked to seasonal affective disorder^15,18,19^ and, later in life, to fragmented activity cycles, cognitive decline, and depression.^20^ The biological benefits of daylight stem from its high intensity and broad spectral composition, which are rarely matched indoors.^21–23^

However, in modern urban life, outdoor time is increasingly limited, and public health interventions to increase daylight exposure require accurate, scalable monitoring. Yet, most studies relied on self-reported diaries and questionnaires, which offer contextual insight but are prone to recall and perceptual biases, often leading to overestimation of outdoor exposure, especially in the context of myopia.^24,25^ Many field studies and health guidelines continue to use simplified proxies, such as the ≥1,000-lux cutoff, to classify and quantify outdoor exposure.^26–29^ While easy to implement, this approach is prone to misclassification as shaded walkways may produce sub-threshold light levels, especially on wrist-worn sensors, even when individuals are technically exposed to outdoor daylight.^21^ Compounding this issue is the limited capability of previously used light sensors, which often record only photopic illuminance or red, green, blue (RGB) values.^30^ These fail to capture the full spectral distribution necessary to evaluate differential stimulation of non-visual photoreceptors, critical for understanding light’s physiological and behavioural effects.^31,32^

Location-based approaches using GPS, Wi-Fi, or inertial sensors^33–35^ offer spatial context but cannot resolve within-location variability related to light exposure—two people only meters apart may receive very different exposures depending on shading, orientation, or window placement.^36^ Moreover, these methods fail to capture the spectral composition of light, limiting their ability to provide biologically and behaviourally meaningful outcomes. This disconnect between measurement and lived experience constrains the relevance of light data for health and behaviour. Bridging this gap requires methods that integrate and compare objective quantification with human perception in real-world contexts. For instance, in our field studies,^37^ participants who were encouraged to spend more time “outdoors” were often unsure what qualified as such.

In this study, we developed *InNOut,* a perception-aligned machine learning framework for classifying and quantifying daylight exposure in everyday life. The model was trained on 73,879 minutes of multispectral light data collected from wrist-worn sensors worn by 22 adults in free-living conditions. Robust ground-truth labels were derived from the sensors’ event button and diary logs. *InNOut* was then benchmarked against independent perceptual judgements of 183 environmental photographs, with concurrent light measurements, labelled as indoor or outdoor by 400 survey respondents, including 17 lighting experts (Fig. 1). This dual-dataset framework moves beyond passive monitoring of light exposure to enable digital behavioural interventions with relatable context-specific recommendations, guiding individuals, workplaces, and urban planners towards environments that meet health-relevant daylight targets, even when full outdoor access is limited.

**Figure 1.**
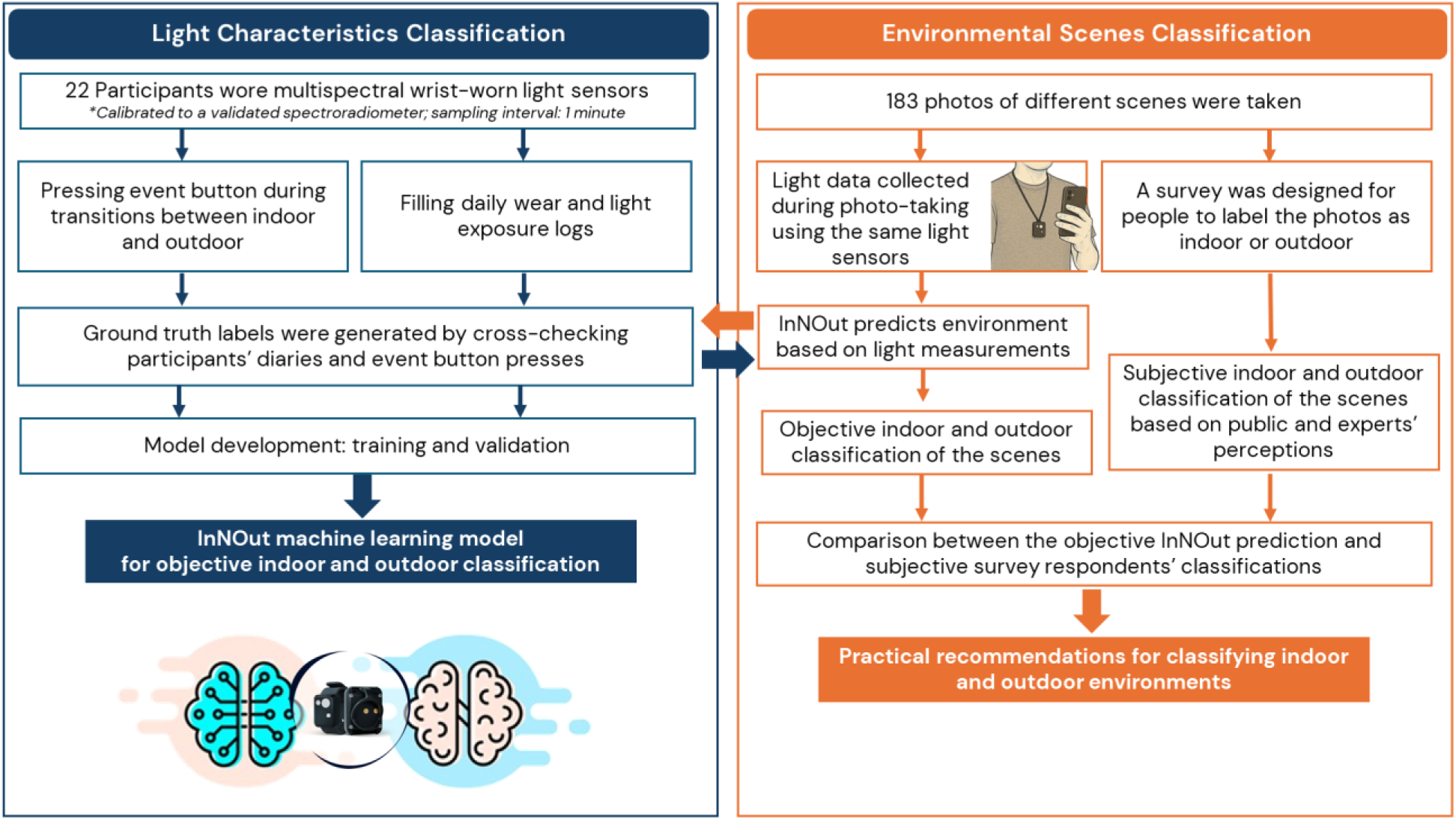
Study design and benchmarking approach. The light characteristics classification (left) involved 22 participants wearing multispectral wrist-worn light sensors for 6.8 ± 0.9 consecutive days in free-living conditions, with ground-truth labels generated from the sensor’s event button presses and daily logs. The environmental scenes classification (right) involved collecting light measurements from 183 photographed scenes and obtaining indoor–outdoor labels from 400 survey respondents, including 17 experts in light and health. *InNOut*’s classifications were compared against human perception to assess agreement.

## RESULTS

We developed and validated *InNOut*, a machine-learning model for classifying indoor versus outdoor environments, using time-series multispectral wrist-worn sensor recordings. The model was first trained and tested on independent datasets of free-living participants (Study 1), and subsequently benchmarked against human perception (Study 2) using a large-scale survey of environmental scenes. Results are presented for model performance metrics (sensitivity, specificity, accuracy, area under the receiver operating characteristics curve (ROC; AUC)), agreement with expert and public perception, and analysis of light-level characteristics across classified environments.

### Study 1: Developing and testing *InNOut*’s performance

#### Time-series multispectral light recordings

Continuous light data were collected from 22 healthy young adults (mean age ± SD = 29.6 ± 4.5 years; 50% male) who wore multispectral wrist-worn sensors (Actlumus, Condor Instruments, São Paulo, Brazil) for an average of 6.8 ± 0.9 consecutive days. Three participants were excluded due to missing or unreliable diaries and/or data, yielding 73,879 minutes of valid data. Robust ground-truth indoor–outdoor labels were generated by cross-checking the sensor’s event button presses with daily light exposure logs. The dataset was split into training/internal testing and external testing sets (Table 1). *InNOut* was trained using gradient boosting machine (GBM) on spectral data and photopic illuminance collected by Actlumus. (more details below). Its performance to classify indoor and outdoor environments in the external testing dataset was compared to the traditionally used 1,000-lux cutoff for photopic illuminance.^21,23,27,28,38–40^

**Table 1:**
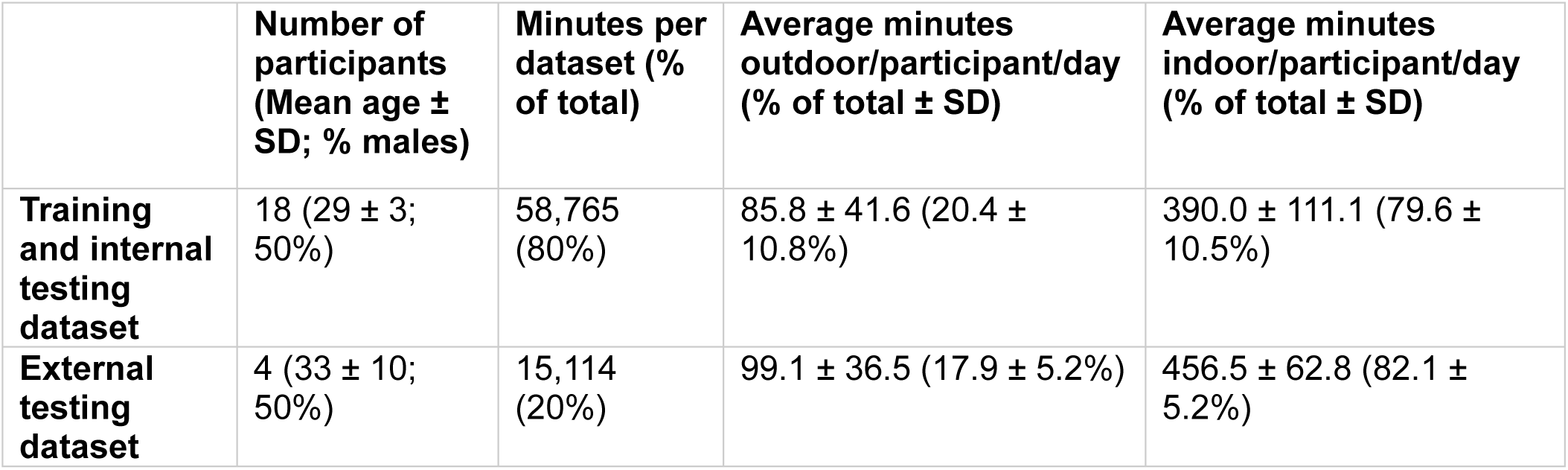
Study 1 dataset characteristics.

#### *InNOut*’s classification performance

In the external-testing dataset, *InNOut* discriminated indoor from outdoor environments with an AUC of 0.92 (95% CI, 0.92–0.93), a sensitivity of 73.9% (72.2–75.5), specificity of 94.5% (94.1–94.9), and an accuracy of 90.9% (90.4–91.3) (Fig. 2A, Supplementary Table 1). *InNOut* achieved a significantly higher AUC compared to the 1,000-lux cutoff (AUC = 0.83 (0.82–0.84); Fig. 2B) in the external testing dataset (DeLong test, *P* < 0.001). *InNOut*‘s sensitivity was higher than the sensitivity of the 1,000-lux cutoff (38.8% (37.0–40.6), t-test with 2000 bootstrap resamples, *P*< 0.001), while maintaining high specificity. Even when matched for specificity (for a 563-lux cutoff), InNOut’s sensitivity remained significantly higher (51.2% (49.3–53.0), P<0.001) than that of photopic illuminance. When data were analyzed per participant, *InNOut* showed consistently high specificity (range: 89.8% to 98.7%) and AUCs (range: 0.87 to 0.97) across participants (Supplementary Table 1).

**Figure 2.**
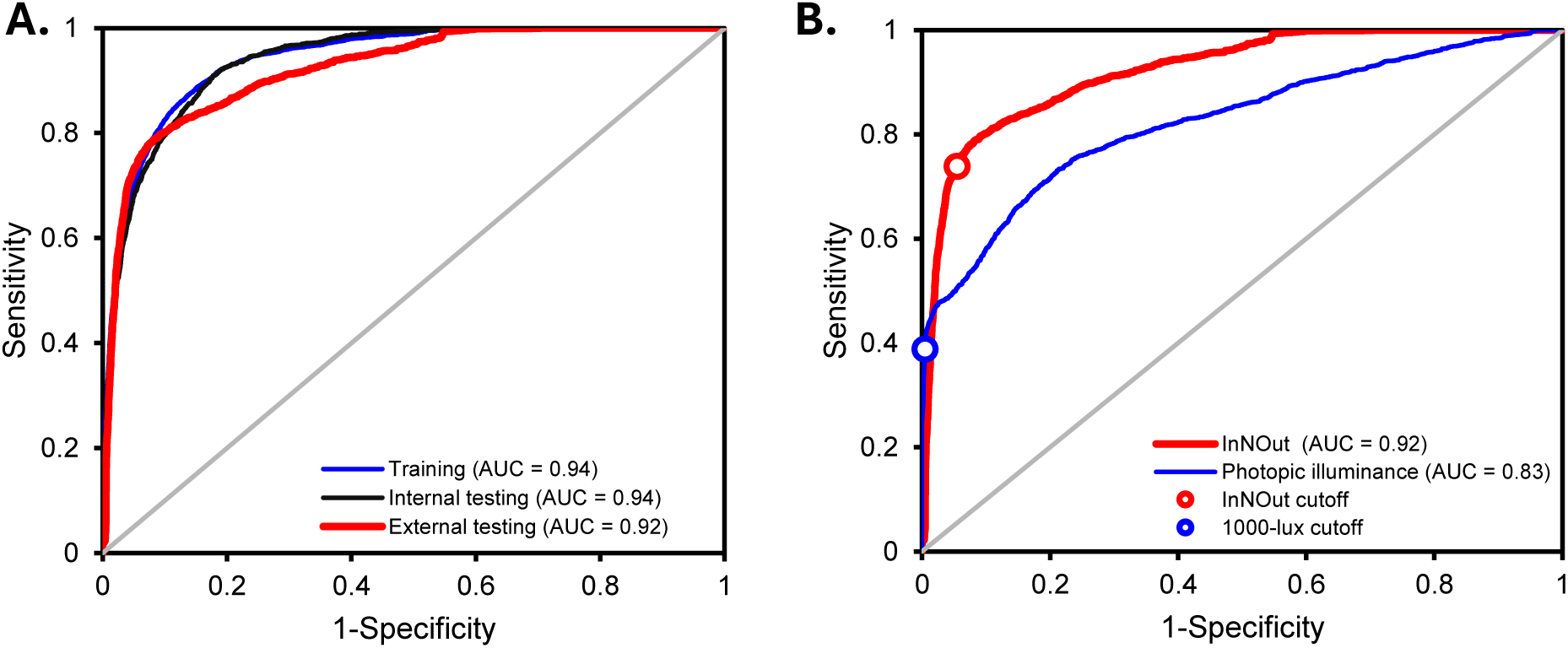
ROC comparing *InNOut* with the 1,000-lux photopic illuminance cutoff for indoor–outdoor classifications. **A.** ROC curves for the training/internal testing and external testing datasets. **B.** *InNOut* (red) outperformed the traditional 1,000-lux cutoff (blue), with higher area under the curve (AUC) values and better discrimination across sensitivity–specificity trade-offs.

#### Consistent Indoor–Outdoor Spectral Differences Across Real-World Measurements

To compare spectral differences between environments, spectral irradiance data were extracted from the raw sensor channels and grouped by environment (indoor vs outdoor). For each sample, a spectral profile was reconstructed directly from the raw channel intensities, and absolute and normalized spectra were then averaged within each environment to obtain representative indoor and outdoor distributions. Across both the Actlumus sensor measurements from participants and the spectroradiometer data, outdoor light consistently showed higher intensities and distinct spectral structure across several wavelength regions (Figure 3A-D). The largest indoor–outdoor differences were observed in the 406–410 nm and 421–425 nm bands, followed by broader differences spanning 471–475 nm, 516–520 nm, 571–575 nm, 661–665 nm, 746–750 nm, as well as the infrared (IR) light channel. These features correspond to the spectral regions that later emerged among the highest-ranked predictors in the GBM model used to train *InNOut*. Photopic illuminance, also captured by Actlumus, ranked fifth in the variable importance of *InNOut* (Fig. 3E).

**Figure 3.**
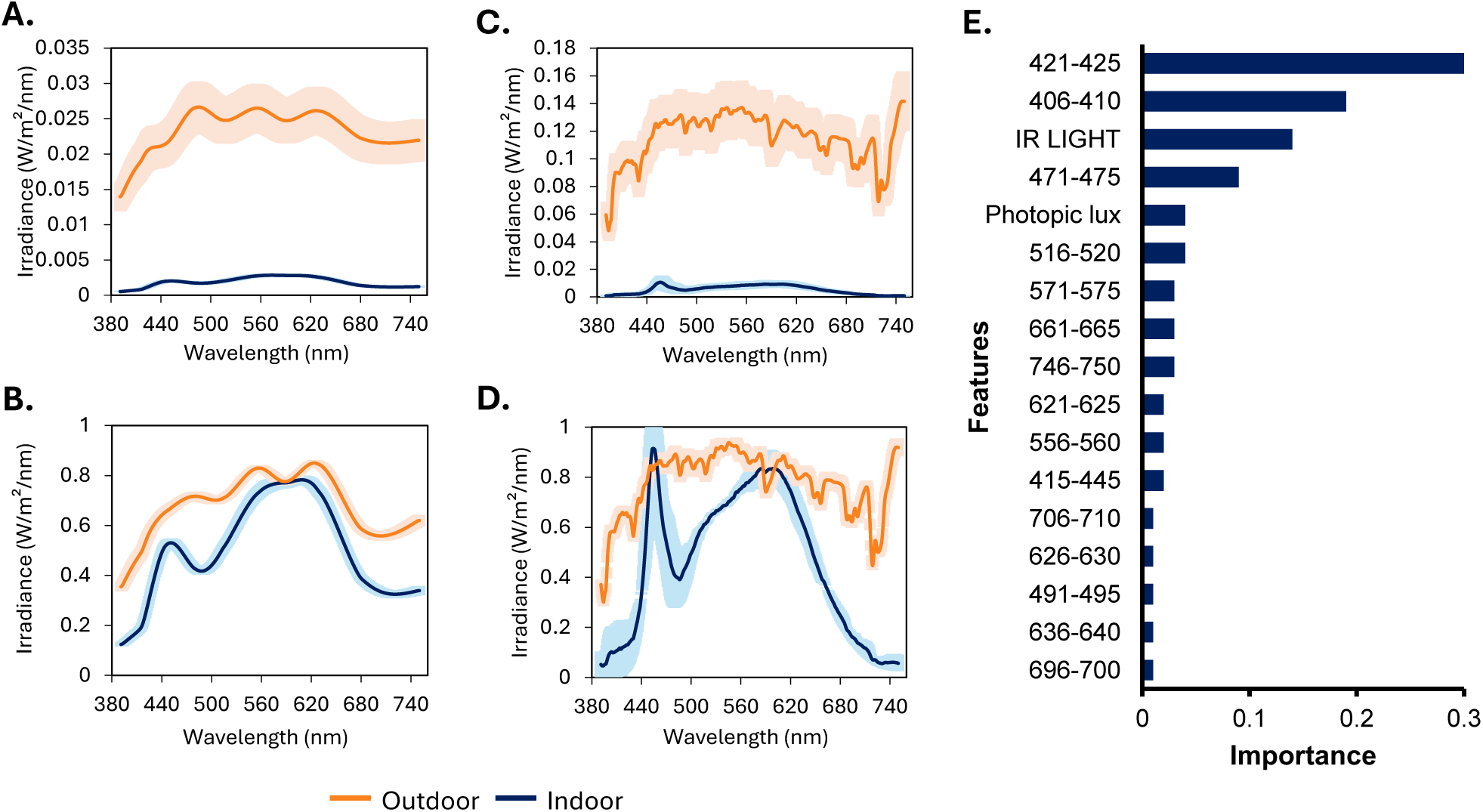
Spectral characteristics of indoor and outdoor environments and feature importance. Averaged participant data from Actlumus light sensors: absolute (A) and normalized (B) spectra. JETI spectroradiometer: absolute (C) and normalized (D) spectra. Feature-importance ranking from the gradient boosting machine model, indicating the relative contribution of individual spectral channels and derived features to indoor–outdoor classification (E).

### Study 2: Environmental scene classification by the public and experts in light and health

In study 2, through an online survey, participants (n = 400, mean age ± SD: 33.9 ± 11.3 years; 43% male), including 17 invited experts in light and health (42.4 ± 7.8 years; 71% male), were required to classify 183 photographs from distinct environments as indoor or outdoor. The spectral lighting characteristics of each photograph were collected using the Actlumus multispectral light sensor, however, this information was not shared with participants. Images were assigned to five main categories and nine subcategories (Table 2).

**Table 2:**
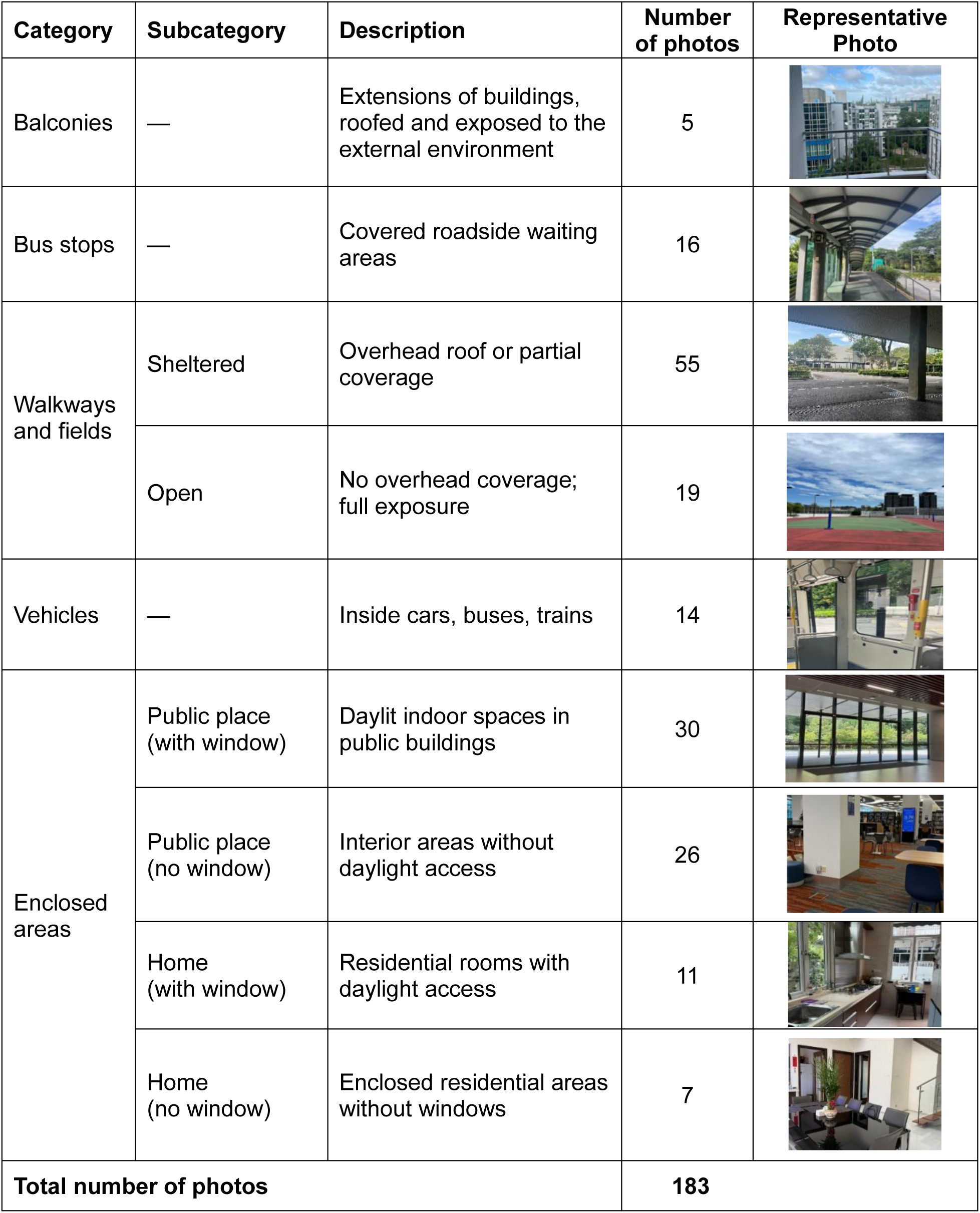
Categories and descriptions of study 2 dataset.

| Category | Subcategory | Description | Number of photos | Representative Photo |
| --- | --- | --- | --- | --- |
| Balconies | — | Extensions of buildings, roofed and exposed to the external environment | 5 |  |
| Bus stops | — | Covered roadside waiting areas | 16 |  |
| Walkways and fields | Sheltered | Overhead roof or partial coverage | 55 |  |
|  | Open | No overhead coverage; full exposure | 19 |  |
| Vehicles | — | Inside cars, buses, trains | 14 |  |
| Enclosed areas | Public place (with window) | Daylit indoor spaces in public buildings | 30 |  |
|  | Public place (no window) | Interior areas without daylight access | 26 |  |
|  | Home (with window) | Residential rooms with daylight access | 11 |  |
|  | Home (no window) | Enclosed residential areas without windows | 7 |  |
| **Total number of photos** |  |  | **183** |  |

#### Consensus patterns in human classification of photographic environments

Of the 183 photographed scenes labelled by survey respondents, 94% met the ≥75% agreement cutoff for indoor–outdoor classification among respondents from the general public (n = 383). Light and health expert respondents (n = 17) exhibited slightly lower overall consistency, with 90% of scenes reaching consensus, diverging particularly in categories where agreement was below the 75% cutoff. These lower-agreement categories were subsequently labelled as “ambiguous” (see methods).

General public respondents classified all bus stops and open walkways/fields as outdoor and all enclosed areas as indoor (100% consensus). Balconies and sheltered walkways/fields were also generally classified as outdoor, with 80% and 87% consensus, respectively, whereas vehicles were classified as indoor in 79% of cases. Experts mirrored these patterns, except for vehicles, for which no consensus was reached. Table 3 summarizes general public and expert consensus rates across environmental categories, highlighting near-complete agreement for most categories and reduced agreement in vehicles and certain semi-open environments.

**Table 3:** Classification rates across environmental categories for experts and general public.

| Environmental Category | % photos in the category reaching consensus* |  | Classification |  |
| --- | --- | --- | --- | --- |
|  | Public | Experts | Public | Experts |
| Balconies | 80 | 80 | Outdoor | Outdoor |
| Bus Stops | 100 | 100 | Outdoor | Outdoor |
| Sheltered walkways and fields | 87 | 82 | Outdoor | Outdoor |
| Open walkways and fields | 100 | 100 | Outdoor | Outdoor |
| Vehicles | 79 | 71 | Indoor | Ambiguous |
| Enclosed areas – Public place (with window) | 100 | 87 | Indoor | Indoor |
| Enclosed areas – Public place (no window) | 100 | 100 | Indoor | Indoor |
| Enclosed areas – Home (with window) | 100 | 100 | Indoor | Indoor |
| Enclosed areas – Home (no window) | 100 | 100 | Indoor | Indoor |

\*Consensus was defined as $\geq 75\%$ agreement within a category, meaning that if at least 75% of the photos in a category received the same label from respondents, that category was assigned that label.

#### Agreement between *InNOut* and the general public and expert perceptions

Light measurements from the photographed scenes were processed by *InNOut* to generate indoor–outdoor predictions for each environmental category. *InNOut*’s classifications closely reflected human perception for most environmental categories, labelling the majority of photographs from open walkways and fields (90%), bus stops (94%), balconies (100%), and sheltered walkways and fields (80%) as outdoors. In contrast, fewer than 20% of photographs from enclosed indoor areas—such as homes and public buildings without windows—were classified as outdoors (Fig. 4). Vehicles (57%) and public indoor spaces with windows (43%) fell within the 25–75% range, indicating perceptually and analytically ambiguous categories according to the cutoff shown in Fig. 4. Overall, *InNOut*’s classifications were in agreement with public perception in 80.8% of cases and expert judgements in 83.5%, with the largest discrepancies occurring in ambiguous categories, which also showed reduced consensus among human raters.

**Figure 4.**
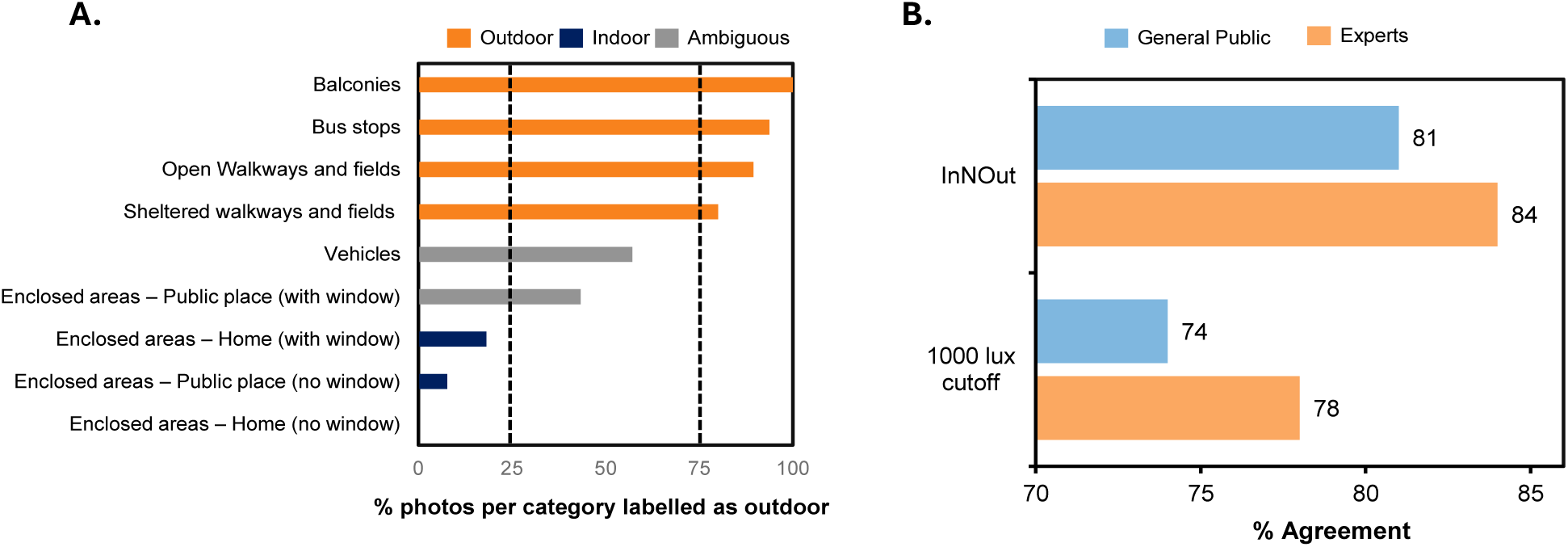
Agreement between *InNOut*’s classifications and human perception across environmental categories. **A.** Bars represent the percentage of photographs in each category classified as outdoor by *InNOut*, with colours indicating model classification outcome: outdoor (orange; ≥75% of photographs classified as outdoor), indoor (blue; ≤25%), and ambiguous (grey; 25–75%). Dotted lines mark the 25% and 75% cutoffsused to define indoor and outdoor categories, respectively. High alignment was observed for open walkways and fields, bus stops, balconies, and sheltered walkways andfields, with lower agreement in more ambiguous categories such as vehicles and public indoor spaces with windows. **B.** Agreement rates between *InNOut* and general public consensus (light blue) and between *InNOut* and expert consensus (light orange) across all scenes.

#### Health-related characteristics of lighting environments

To assess the health-related characteristics of light exposure across different environments, we analysed photopic illuminance and melanopic equivalent daylight illuminance (mEDI) values measured in the photographed scenes. Measured light levels varied systematically across photographic categories and were consistent with the model’s indoor–outdoor classifications. Open and semi-open environments—including balconies, bus stops, and vehicles—exceeded both the commonly used photopic illuminance threshold for outdoor classification for myopia research (≥1,000 lux)^21,23,27,28,38–40^ and the mEDI threshold associated with non-visual benefits (≥250 lux)^41^.

In contrast, fully enclosed settings, particularly those without windows, consistently fell below both thresholds. Public indoor spaces with windows showed intermediate values, reflecting their position as perceptually and photometrically ambiguous environments (Fig. 5, Supplementary Fig. 1).

**Figure 5.**
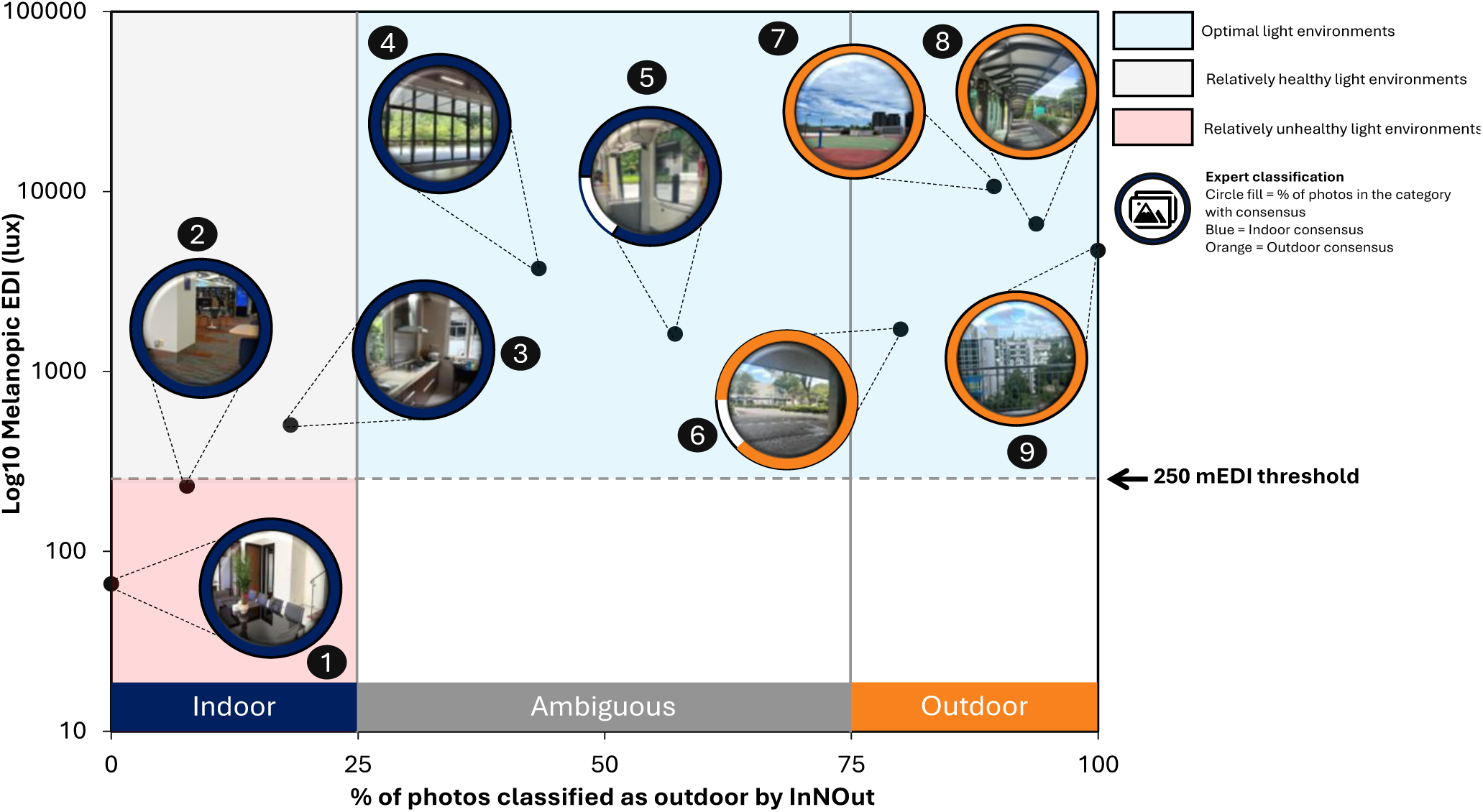
Integration of *InNOut*, expert consensus, and healthy light exposure across nine environment categories. The x-axis shows the percentage of photographs classified as outdoor by the *InNOut* model within the environment category, with three zones: indoor (blue; ≤25%), ambiguous (grey; 25–75%), and outdoor (orange; ≥75%). The y-axis shows actual light levels as melanopic EDI (lux, log₁₀ scale). Background shading indicates health relevance: blue = optimal light environments (≥250 mEDI, outdoor or ambiguous), grey = relatively healthy light environments (>250 mEDI, indoor, next to a window), and red = relatively unhealthy light environments (<250 mEDI, indoor). The dashed line marks the 250 mEDI threshold for beneficial daytime light exposure. Expert classifications are represented by the ring color (blue = indoor consensus, orange = outdoor consensus), while circle fill indicates the percentage of photographs within each category for which experts reached consensus. Values between 1 and 10 log10 mEDI were not represented in this plot given that all photographs were taken during daytime. **Categories:** (1) Enclosed Area – Home, (2) Enclosed Area – Public Place, (3) Enclosed Area – Home (window), (4) Enclosed Area – Public Place (window), (5) Vehicle, (6) Sheltered Walkways/Spaces, (7) Open Walkways/Spaces, (8) Bus Stops, (9) Balconies.

## DISCUSSION

Our findings demonstrate that *InNOut* bridges a long-standing gap between objective light measurements and subjective human perception in real-world environments. By aligning spectral data with how people perceive indoor and outdoor contexts, *InNOut* offers a biologically meaningful yet intuitively relatable framework for classifying light exposure and provides a more context-aware approach to quantifying light environments relevant to health, an advance over cutoff/threshold-based^26–29^ and location-tracking^33–35^ methods that often misrepresent lived experience. In external testing, *InNOut* achieved high performance (AUC: 0.92; accuracy: 90.9%; sensitivity: 73.9%; specificity: 94.5%), with 35% improvement in the detection of true outdoor exposure (i.e., sensitivity) compared to the traditionally adopted 1,000-lux cutoff for outdoor classification. This improvement is particularly important as some outdoor environments, such as shaded areas between buildings or under shelters, may fall below the illuminance threshold despite having the spectral properties of daylight and potentially being health-promoting, and some indoor spaces with large windows may exceed it.^21,42^ These findings highlight the limitations of relying solely on photopic illuminance, commonly used in previous research,^21,23,27,28,38–40^ but increasingly questioned as an inadequate metric,^43^ and emphasize the value of context-aware classification for understanding light exposure in real-world settings. Integrating *InNOut* into digital behaviour-change programs would allow the correction of misconceptions through objective, data-driven feedback, guiding individuals toward environments that reliably deliver health-relevant light exposure. In doing so, the framework extends beyond accurate classification to serve as a tool for sustainable, data-driven behaviour change. With larger, multi-center datasets, it will also enable country-specific insights, allowing researchers to move beyond general recommendations to increase outdoor time and instead offer clear, contextually relevant definitions of “outdoors” to study participants.

Validation against human classification further demonstrated the model’s interpretability and the study’s human-centered design. Prior studies have highlighted a disconnect between objective light metrics and subjective light perception.^24,25,38,44^ Our approach addresses this gap by linking spectral data to real-world photographic scenes, producing recommendations that align with how people perceive their surroundings. Across 183 photographs, *InNOut* matched public perception in 80.8% of cases and expert consensus in 83.5%. Human consensus showed that open walkways, bus stops, and balconies were almost universally perceived as outdoor, while enclosed spaces without windows were consistently classified as indoors. Ambiguous spaces such as vehicles and partially windowed public interiors showed greater variability in perception, and *InNOut* mirrored this uncertainty, reflecting the genuine complexity of these settings^45,46^. This strong perceptual alignment enables objective sensor data to be translated into contextual guidance that individuals recognize and trust. When paired with photographs of environments that reliably provide—or fail to provide—healthy light, the model’s outputs become actionable, supporting behaviour-changing applications, personalized recommendations, and public health guidance that remain scalable, privacy-preserving, data-driven, and relatable^47^ to urban populations with limited daylight access.

Furthermore, light-level analysis confirmed that outdoor and semi-outdoor spaces, including balconies, open fields, and sheltered walkways, consistently exceeded thresholds recommended for health-relevant exposure (≥250 melanopic lux for daytime),^41^ while enclosed indoor spaces without windows fell well below these targets. These patterns have direct practical implications. For individuals unable to spend prolonged periods outdoors, semi-open spaces offer a viable alternative for achieving health-relevant light exposure. Public health and workplace guidelines could therefore encourage breaks in such settings, and architectural design can maximize daylight access in public buildings. It is important to note, however, that although public places with windows and vehicles were classified as ambiguous in our classification framework, their potential protective effects with respect to myopia prevention remain unconfirmed and warrant further investigation.

Critically, *InNOut* enables these recommendations to move beyond abstract guidance into actionable, data-driven behaviour change. Embedded in wearable platforms, the model can provide individuals with real-time information on whether their environments deliver health-relevant light. This objective feedback corrects common misconceptions, such as counting malls^48^ or dimly lit interiors as outdoor time and helps people adapt routines in ways that truly support circadian and ocular health. At the programmatic level, workplaces, schools, and clinicians could incorporate *InNOut*-based feedback into wellness initiatives, allowing progress to be tracked, goals set, and interventions adjusted dynamically. In this way, *InNOut* transforms light exposure monitoring from a descriptive tool into a digital behaviour-changing framework that empowers both personal decision-making and population-level health strategies.

The current work has limitations that also serve as opportunities for future development. The model was developed and validated using a single wearable light sensor (Actlumus), which may require adaptation for other devices. Future work should explore cross-sensor calibration methods or open-source spectral response profiles to facilitate interoperability. The current device lacks UV sensing, which could be added to extend applications to contexts where ultra-violet (UV) exposure is relevant. Data collection was limited to Singapore, and expanding to other geographies, seasons, and architectural styles will allow assessment of generalizability. Ground-truth labels were derived from diaries and event button presses, which are prone to human error and recall bias. While cross-validation and external testing were employed to mitigate these issues, future work should explore automated annotation methods to improve label fidelity. While these limitations define the current scope of our work, they also provide a roadmap for expansion. Applying *InNOut* across diverse devices, geographic regions, seasons, and architectural contexts will be critical for testing generalizability, while integrating temporal light dynamics and additional environmental cues may improve classification in perceptually ambiguous settings.

In this study, we offer a practical pathway from measurement to intervention by identifying which everyday settings reliably deliver health-relevant light exposure and translating these findings into concrete, context-specific recommendations, such as using balconies or covered walkways during breaks, favoring open over enclosed routes, or seeking indoor spaces with generous daylight access when outdoor time is limited. This perception-aligned framework can inform personal behaviour, workplace wellness strategies, and urban design, moving light monitoring beyond passive quantification toward active, context-aware guidance that supports ocular health, circadian stability, mood regulation, and broader wellbeing.

## METHODS

### 1. Data Collection

#### Study 1: Developing and testing *InNOut*’s performance

##### Participants

Twenty-five healthy adults (mean age ± SD = 29.5 ± 4.2 years; 48% male) wore Actlumus sensors on the non-dominant wrist for 6.8 ± 0.9 consecutive days covering all daily activities except showering and water-related activities. Participants were eligible if they were aged 21–45 years, able to commit to the wear protocol, and provided written informed consent. Participants pressed the sensor’s event button when transitioning between indoor and outdoor environments and maintained diary light-exposure logs. Ground-truth labels were established by cross-referencing button-press timestamps with diary entries using predefined reconciliation rules (e.g., if entries disagreed, the event was excluded to avoid mislabelling). Non-wear periods (detected by the device’s off-wrist sensor and cleaned based on inactivity) and light data outside 07:00–19:00 local time were excluded. Three participants were removed due to missing/unreliable diaries or data, yielding a final sample of 22 participants (mean age ± SD = 29.6 ± 4.5 years; 50% male) and 73,879 minutes of valid data.

##### Ambulatory multispectral light recording

Environmental light exposure was measured using Actlumus wrist-worn multispectral sensors (Condor Instruments, São Paulo, Brazil), which capture light across 10 spectral channels spanning violet to infrared (measurement range 1–100,000 lux). Devices also record activity and skin temperature via embedded accelerometers and thermistors. Sensors were worn on the non-dominant wrist and sampling was configured at 60-second intervals. Actlumus devices were calibrated against a reference spectroradiometer (JETI Spectraval 1511, Technische Instrumente GmbH, Germany). A comparison between indoor and outdoor measurements using both devices is presented in Fig. 3A-D.

##### Development of the gradient boosting classification model (*InNOut*)

*InNOut* was developed using a gradient boosting machine (GBM) applied to time-series multi-channel light recordings collected in Study 1. Each timepoint comprised 10 raw spectral channel values covering the visible wavelength range (390–750 nm). Recordings were labelled according to the experimental condition, with *EVENT = 0* denoting indoor and *EVENT = 1* denoting outdoor environments. For each observation, the spectral intensity profile was reconstructed directly from the raw wavelength-specific channel measurements by interpolating the spectral bands across the 390-750 nm range. These spectral profiles were then averaged across all indoor and outdoor samples separately, yielding representative mean spectral distributions for each condition. Spectral features showing the largest differences between environments (e.g., 406–410 nm, 421–425 nm, etc.), along with photopic illuminance values, were selected as model inputs. When two features were highly correlated (*R* > 0.8), only the one with the greater variable importance was retained. A 15-minute sliding window (15 consecutive 1-minute samples) was applied to enhance temporal stability and reduce noise. Data from 18 participants (58,765 minutes; 80%) were used for model training and cross-validation, while data from 4 participants (15,114 minutes; 20%) served as an external test set. For comparison, a threshold-based classifier using a 1,000-lux photopic illuminance cutoff was evaluated on the same dataset. All models were implemented in Python 3.9.13 using the *GradientBoostingClassifier* function from the *scikit-learn* library (version 1.6.1).

#### Study 2: Environmental scene classification by the general public and experts in light and health

A database of 183 geo-tagged photographs representing everyday environments (e.g., public places, residential areas, vehicles) all around Singapore was assembled. During each photo capture, a team member wore the Actlumus sensor on a chest-mounted lanyard to record concurrent multispectral light measurements and key contextual variables (*e.g.,* time of day, weather, presence of windows) were manually noted. Photos captured heterogeneous settings with varying degrees of exposure and enclosure and were grouped post hoc into environmental categories based on built features (*e.g.,* roofing, window presence). The image set formed the basis for a classification survey delivered via Microsoft Forms to two groups: (i) adults from the general population (age ≥ 21 years) and (ii) selected invited experts (PhD holders with ≥3 years’ experience in myopia research, circadian biology, photobiology, architecture, or lighting engineering). For each photograph, respondents classified the scene as either “indoor” or “outdoor.”

##### *InNOut* for classification of photographic environments

Spectral data recorded during the capture of the photographs were processed through the trained *InNOut* model. To maintain consistency with the training set, only photographs taken between 07:00 and 19:00 local time were included. A total of 183 photographs met quality-control criteria (valid spectral data, complete contextual records) and were classified as “indoor” or “outdoor.” Model predictions were then compared with human survey classifications to assess perceptual alignment and evaluate the model’s generalisability across novel environmental contexts.

## Statistical analysis

Model performance was assessed using ROC curve analyses. Classification metrics included accuracy, sensitivity, specificity, and AUC. Bootstrapping (2,000 resamples) was used to compute 95% confidence intervals for each metric. AUCs were compared using DeLong’s test to evaluate performance differences between *InNOut* and the traditional 1,000-lux cutoff.

Survey responses were analysed using a consensus threshold of 75%, whereby a photograph was assigned an “indoor” or “outdoor” label if ≥75% of respondents agreed, following precedent for consensus thresholds in other studies^49,50^ For categorical analysis (e.g., balconies, sheltered walkways), a label was assigned to the category if ≥75% of the photographs within it shared the same classification; otherwise, the category was considered ambiguous.

## Author contributions

Concept and design: DM and RPN. Data acquisition: DM. Analysis and interpretation: DM, ZT, and RPN. Manuscript preparation: DM and RPN.

## Acknowledgments

We would like to thank the research participants, survey respondents, and the students that helped with photo taking for the survey development: Casey Huang, Raphael Ho Jun Wei, Sophia Marie Bagui Dela Cruz. This research was funded by the ASPIRE-NUS startup grant (NUHSRO/2022/038/Startup/08) to RPN.

## Data availability

The de-identified datasets used in this study can be made available from the authors upon reasonable request. The full study protocol is available from the corresponding author on reasonable request.

## Code availability

The code used in this study can be made available from the corresponding author upon reasonable request.

## Ethical approval and consent for publication

Consent was obtained directly from participants. Both studies were approved by the National University of Singapore Institutional Review Board (NUS IRB), study 1: NUS-IRB-2023-863; study 2: NUS-IRB-2024-1023.

