## Supplementary Material for "Human-Perception-Aligned Machine Learning for Indoor–Outdoor Classification"

**Corresponding Author:**

Dr Raymond P. Najjar

### This supplementary material document includes:

**Supplementary Table 1.** Classification performance of *InNOut* compared with the 1,000-lux cutoff

**Supplementary Figure 1.** Melanopic equivalent daylight illuminance (mEDI) across environmental categories.

**Supplementary Table 1. Classification performance of *InNOut* compared with the 1,000-lux cutoff**

| Source | Sensitivity (%) | Specificity (%) | Accuracy (%) | AUC (95% CI) |
|---|---|---|---|---|
| **Training** | 65.9 (64.8–66.9) | 95.8 (95.6–96.0) | 90.7 (90.4–90.9) | 0.94 (0.94–0.94) |
| **Internal testing** | 64.7 (62.7–66.9) | 95.6 (95.2–96.0) | 90.3 (89.8–90.9) | 0.94 (0.94–0.94) |
| **External testing** | **73.9 (72.2–75.5) ***** | **94.5 (94.1–94.9) ***** | **90.9 (90.4–91.3) ***** | **0.92 (0.92–0.93) ***** |
| LL06 | 70.1 (67.4–72.6) | 98.7 (98.3–99.0) | 91.8 (91.0–92.5) | 0.97 (0.97–0.97) |
| LL11 | 76.6 (73.3–79.8) | 95.1 (94.4–95.7) | 92.7 (92.0–93.4) | 0.95 (0.94–0.95) |
| LL17 | 69.9 (66.0–73.8) | 91.3 (90.3–92.3) | 88.1 (87.1–89.2) | 0.87 (0.85–0.89) |
| LL21 | 86.8 (83.1–90.2) | 89.8 (88.3–91.2) | 89.2 (87.9–90.6) | 0.93 (0.91–0.94) |
| **1,000-lux cutoff** | **38.8 (37.0–40.6)** | **99.5 (99.3–99.6)** | **88.6 (88.1–89.1)** | **0.83 (0.82–0.84)** |

*Performance metrics (95% confidence intervals) for training, internal testing, and external-testing datasets, and for individual participants in the external testing set (Participants LL06, LL11, LL17, LL21). Metrics include sensitivity, specificity, accuracy, and area under the ROC curve (AUC). *** P < 0.001 when comparing InNOut to the 1,000-lux cutoff.*

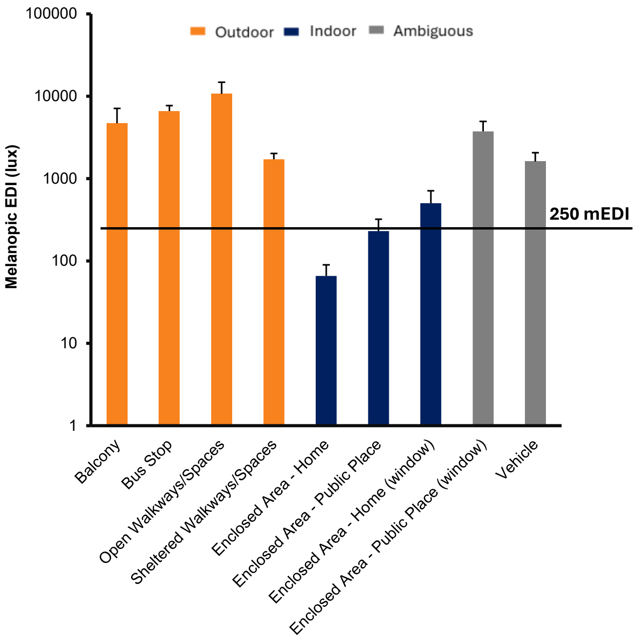


**Supplementary Figure 1. Melanopic equivalent daylight illuminance (mEDI) across environmental categories.** Bars represent mean values for each category, with colours indicating *InNOut* classification outcome: outdoor (orange; ≥75% of photographs classified as outdoor), indoor (blue; ≤25%), and ambiguous (grey; 25–75%). The y-axis shows actual light levels as melanopic EDI (lux, $\log_{10}$ scale). The black line marks the ≥250 mEDI threshold associated with health benefits. Open and semi-open environments typically exceeded the threshold, while fully enclosed settings, particularly those without windows, fell below it. Error bars indicate the standard error (SE).
